# Identification of multi-targeting and synergistic neuromodulators of epilepsy associated protein-targets in Ayurvedic herbs using network pharmacological approach

**DOI:** 10.1101/471474

**Authors:** Neha Choudhary, Vikram Singh

## Abstract

Epilepsy comprises a wide spectrum of neuronal disorders and accounts for about one percent of global disease burden affecting people of all age groups. In the traditional medicinal system of Indian antiquity, commonly known as Ayurveda, epilepsy is recognised as *apasmara* and a plenty of information is documented regarding the effectiveness of various herbs against it. Towards exploring the complex regulatory mechanisms of Ayurvedic herbs at molecular levels, in this study, a network pharmacological framework is developed for thoroughly examining the anti-epileptic potential of 349 drug-like phytochemicals (DPCs) found in 63Ayurvedic herbs. Interaction networks of phytochemicals in anti-epileptic herbs, their protein targets and associated human pathways are designed at various scales and DPCs are mapped on these networks to uncover complex interrelationships. Neuromodulatory prospects of anti-epileptic herbs are probed and, as a special case study, DPCs that can regulate metabotrophic glutamate receptors (mGluRs) are inspected. An account of novel regulatory phytochemicals against epilepsy is reported by systematically analysing the screened DPCs against DrugBank compounds. A repertoire of DPCs having poly-pharmacological similarity with anti-epileptic drugs available in DrugBank and those under clinical trials is also reported. Further, a high-confidence PPI network specific to the protein targets of epilepsy was developed and the potential of DPCs to regulate its functional modules was investigated. The study concludes by highlighting a couple of herbs as potential sources of epileptogenic regulators. We believe that the presented schema can open-up the exhaustive explorations of indigenous herbs towards meticulous identification of DPCs against various diseases and disorders.

## 1. Introduction

Epilepsy (EP) is one of the world’s oldest known conditions of neuronal disorder, with a history dating back to 4,000 BC of misconception, unawareness and humiliation (Valeta, 2017). This neurological disorder has been one of the most researched medical condition mainly due to the complex morbidity associations with substantially high mortality rates (Mohanraj et al., 2006; Yuen et al., 2007). High mortality is considered to be the consequence of underlying disorders associated with the disease that may include non-epilepsy related other factors also (Tomson, 2000; Nevalainen et al., 2014). EP is amongst the top three leading contributors to global burden of neurological disorders, affecting about 65 million people worldwide with widely varying prevalence and incidence throughout the world (Devinsky et al., 2018). Earlier reported studies have shown that epilepsy-related mortality is 2.6 fold higher in low and middle-income countries than the general population, where 80-90% of epilepsy patients receive no treatment at all (Levira et al., 2017; WHO, IBE and Atlas, 2005). The prevalence of this disorder in South East Asia is 2-10 persons per 1000 population of which more than 50% of DALYs (Disability-adjusted life years) are contributed from India. Further, the effect of this disorder on children is quite disturbing. Population-based studies have shown that the risk of death is 10 times higher in case of children, compared to the general population (Donner et al., 2017). For the childhood epilepsy, complex interactions among underlying epileptic co-morbidities emerging in the developing young brain constitute the major treatment challenge (Wilmshurst et al., 2014).

EP is a chronic neuronal disorder of the brain that is characterized by recurrent and spontaneous seizures (epileptic seizures) leading to events of involuntary movement of the body or body parts. An epileptic seizure is a clinical manifestation accredited to the synchronous neuronal activity within the brain (Fisher et al., 2005). Based on the aetiology, EP can be categorized into two categories: symptomatic and idiopathic EP. While symptomatic EP could be remote (conditions attributed to head injury, stroke, CNS infection) or progressive (brain tumor and degenerative disease) in nature, idiopathic has no identifiable cause. It should be taken into consideration that the complex behavior of EP is also attributed to the psychological, social and economic backgrounds (Devinsky et al., 2018). Although remarkable advancements towards the understanding and treatment of EP have been achieved in past, existence of neurological and psychiatric co-morbidities undermine the current treatment strategies of EP (Kanner, 2016). Further, the conventional approach of “one drug-one target-one disease” treatment schema seems to be unsuccessful in providing satisfactory results. Since the locus of the seizure in the majority of EP patients is not confined to a single region of the cortex, even surgery is not opted as a treatment strategy in such cases. Also, the use of new generation antiepileptic drugs are less successful in completely controlling the seizures, thus providing a scope for improvements in current drug-development procedures (Löscher et al., 2013). Drug-resistant seizures have also been observed in nearly one-third of the epileptic patients (Kwan and Brodie, 2000). Considering all these factors into account, network medicine approach for the treatment of EP could be of significant importance. Owing to their suitability in the treatment of complex diseases including drug-resistant cases, ideas that can integrate the concepts of network biology with polypharmacology are drawing attention of the scientific community (Hopkins, 2008).

Since herbal medicines that are one of the most popular forms of CAMs (complementary and alternative medicines) are effective and safe in nature, their usage in the treatment of neurological diseases and disorders also is gaining importance. Herbal formulations exert their poly-pharmacological effects by acting on the multiple targets using its multi-component framework (Wu et al., 2013). Trend of the treatment of EP using CAMs is not new and nearly 50% of the patients are still using this strategy for the treatment (Ricotti and Delanty, 2006). Ayurveda, the ancient Indian medicinal system defines EP as *Apasmara:* where *apa* refers to “negation” and *smara* as “consciousness” (Manyam, 1992). Ayurveda describes the use of various plants and plant-based formulations for the management of neurological conditions like migraine, epilepsy, Alzheimer’s, anxiety, Parkinson’s, depression etc. (Ayurvedic Pharmacopoeia of India, 2001; Ayurvedic Formulary of India, 2000). Due to rapid advancements in the field of chemical and synthetic biology, the network pharmacological paradigm of drug discovery is showing potential to be efficient and effective in exploring the molecular mechanisms of various herbs like *Piper longum* (Choudhary and Singh, 2018), herbal formulae’s like Yu Ping Feng decoction (Zuo et al., 2018) and therapeutic molecular classes like cannabidiol (Bian et al., 2018) etc. It is a poly-pharmacological approach which relies on the principle of “multiple compounds – multiple-targets” approach for the disease treatment and represents the holistic mechanism of the drugs, targets and disease in a systematic manner.

Therefore, in the present study, we have first identified the extensively used anti-epileptic herbs of Ayurveda and then their poly-pharmacological evaluation was carried out using the approach of network pharmacology. For this, we have manually reviewed and identified the anti-epileptic herbs of Ayurveda from various literature resources. A comprehensive dataset of phytochemicals present in identified herbs was prepared using the information present in public databases and was analyzed for their drug-likeliness properties and chemical classification. Further, the information of protein molecules targeted by phytochemicals was collected and poly-pharmacological action of each was assessed based on the number of protein targets that a phytochemical may target. Functional significance of protein targets was assessed using the pathway and module associations. The high confidence phytochemical – protein target pairs involved in this disorder were identified and their molecular interactions were assessed on the basis of *in-silico* docking studies. The complete methodology of this study is presented in Fig. 1.

**Figure 1.**
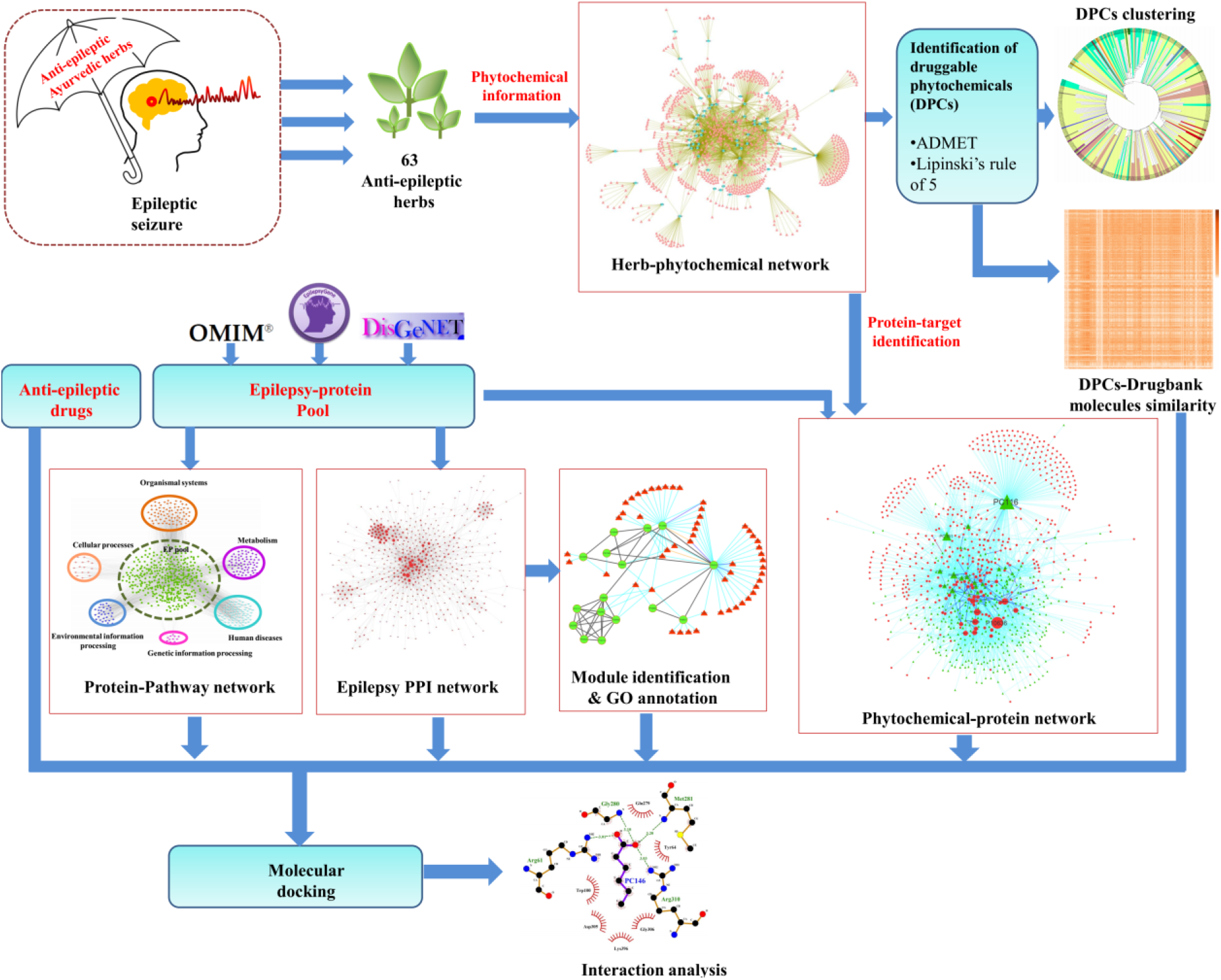
Detailed workflow of the present study.

## 2. Material and methods

### 2.1. Identification of anti-epileptic herbs (AEHs) and drugs (AEDs)

To collect the information of the anti-EP herbs prescribed in Ayurveda system of medicine, relevant research articles present in PubMed-NCBI (https://www.ncbi.nlm.nih.gov/pubmed/) were selected and manually inspected. For the identification of the AEDS that are in current use and also under clinical trials, DrugBank database (https://www.drugbank.ca/) (Wishart et al., 2018) screening and literature survey were performed.

### 2.2. Phytochemical dataset preparation of anti-epileptic herbs

The list of phytochemicals present in the identified Anti-epileptic herbs was compiled from 3 database sources: Duke’s phytochemical database (https://phytochem.nal.usda.gov/phytochem/search), TCMSP (Traditional Chinese Medicine Systems Pharmacology) (Ru et al., 2014), and PCIDB (PhytoChemical Interactions DB) (https://www.genome.jp/db/pcidb). PubChem (https://pubchem.ncbi.nlm.nih.gov/) and ChEMBL (https://www.ebi.ac.uk/chembl/) databases of chemical compounds were used to derive the chemical information of phytochemicals. 2D and 3D chemical structures of phytochemicals were obtained using OpenBabel 2.4.1 software (O’Boyle et al., 2011). The phytochemicals, for which no chemical information was present, are not considered in this study.

### 2.3. Protein target identification of AEDs and phytochemicals in AEHs (phytochemical dataset)

The identification of potential drug targets is considered a key step toward the drug development procedure (Schenone et al., 2013; Hughes et al., 2011). Here, three target prediction software was used to collect the information of the potential protein targets of the phytochemical dataset. (1) BindingDB (https://www.bindingdb.org/), a web-accessible public database that contains the protein interaction information of the potential drug targets with their ligand molecules (Liu et al., 2007). To access the high confidence targets, BindingDB was searched with interaction score of ≥ 0.85. (2) STITCH 5.0 (http://stitch.embl.de/), which catalogs the information of the manually curated as well as experimentally validated protein-chemical interactions (Szklarczyk et al., 2016). Only the chemical interaction with the STITCH confidence score of ≥ 0.4 was considered. (3) Swiss Target Prediction (http://www.swisstargetprediction.ch/), a web server for the target identification of bioactive small molecules that uses the combination of 2D and 3D similarity measures for the prediction of top-15 targets (Gfeller et al., 2014). For all the predictions, the search was limited to “*Homo sapiens*”.

### 2.4. EP targets (EP-gene pool)

The data of EP associated genes were collected from three databases: (1) EpilepsyGene (http://www.wzgenomics.cn/EpilepsyGene/), a genetic resource which accounts for the information on epilepsy-related genes and mutations, collected from various research publication (Ran et al., 2015); (2) OMIM database (http://omim.org/), a continuously updated catalog of human genes and genetic phenotypes establishing a relationship between phenotype and genotype; and (3) Disgenet, a collection of disease-associated genes and variants associated with *Homo sapiens* (Piñero et al., 2017). The list of 1179 genes corresponding to epilepsy is listed in supplementary file1, Table S1.

### 2.5. Compound classification and clustering

The chemical classification of phytochemicals was obtained using “Classyfire”. This provides the hierarchical classification of chemical compounds based on the chemical ontology of approx. 4825 organic and inorganic compounds (Djoumbou Feunang et al., 2016). Clustering of the compounds based on atom-pairs descriptors and Tanimoto coefficient were performed using ChemMine tools (Backman et al., 2011).

### 2.6. Pharmacokinetic prediction and Drug-likeliness evaluation

Estimation of pharmacological properties of small molecules is considered a crucial step towards the drug discovery approaches. In that direction, owing to the disadvantage of *in vivo* estimation as time consuming and expensive, *in silico* methods have become inevitable approaches. In this study, pkCSM, a graph-based signature method was used to predict the pharmacokinetic and toxicity properties of the compounds (Pires et al., 2015). Four ADMET related parameters including Lipinski “rule of five” criterion, caco2 permeability, intestinal absorption and toxicity assessment were used to screen drug-like molecules from the list. Lipinski’s “rule of five” criterion is considered as a basis for estimating the drug like nature of the chemical compounds. According to its five component measure, a chemical compound must have H-bond acceptors less than 10, number of H-bond donors less than 5, molecular weight less than 500 Dalton and logP value of less than 5 for its proper absorption and permeability (Lipinski et al., 1997). Further, the permeability coefficient across the human colon carcinoma cell lines caco2 is also considered as an important parameter to predict the absorption of the orally administered drugs (Hubatsch et al., 2007). For this, the threshold of caco2 permeability score > 0.9 and intestinal absorption value of > 30% was considered as a cut-off. Lastly, phytochemicals negative for Hepatotoxicity and Ames toxicity were selected. Only phytochemicals passing these above criteria were selected for subsequent analysis and declared as putative drug-like molecules. The ADMET (Absorption, Distribution, Metabolism, Excretion and Toxicity) properties of 349 putative drugable phytochemicals referred to as DPCs are provided in supplementary file2, Table S2.

### 2.7. Similarity index calculation

For evaluating the similarity between the 349 DPCs and the existing drugs available at DrugBank, Tanimoto coefficient (TC) was calculated using OpenBabel (version 2.4.1) (O’Boyle et al., 2011). A path-based fingerprint which indexes small molecule fragments upto the length of seven atoms *i.e*. FP2 was used for the TC calculation. TC between chemical compound C_1_ and C_2_ is given by,

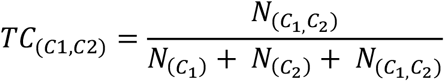

where, *N*_(*C*_1_)_ and *N*_(*C*_2_)_ referes to the number of molecular fingerprints present in C_1_ and C_2_ respectively. While *N*_(*C*_1_,*C*_2_)_ represents number of molecular fingerprints common to both C_1_ and C_2_. Range of the TC varies from 0-1, where 0 represents the minimun while 1 represents the maximum similarity.

Only the drugs corresponding to “drugbank_approved_structure” data, present in DrugBank dataset library were selected for the comparison. The Tanimoto score of each of 349 DPCs and 2159 drugs are given in the supplementary file3, Table S3.

### 2.8. Network analysis

The PPI (protein-protein-interaction) data of protein targets was fished from STRING, with search limited to “*Homo sapiens*” and confidence score ≥ 0.9 (Szklarczyk et al., 2015). Cytoscape (version 3.0), an open source software was used for the network construction, analysis and visualization (Shannon et al., 2003). Network construction and analyses were performed for the following cases: (1) Herb-Phytochemical (H-PC) network, (2) Druggable phytochemical-Protein target (DPC-PT) network, (3) Protein target-Human pathway (PT-HP) network, and (4) Epilepsy PPI (EP-PPI) network. The pathway database of KEGG (Kyoto Encyclopedia of Genes and Genomes, (http://www.genome.jp/kegg/pathway.html) was used to retrieve the information of human pathways associated with the protein targets (Kanehisa et al., 2012). Molecular Complex Detection (MCODE) algorithm was opted to identify and organize the functional modules in the EP-PPI network. These densely connected regions *i.e*. modules are known to possess network like properties and tend to be composed of a set of proteins that in coordination with each other regulate specific functions (Barabási et al., 2011). Clusters were analyzed for the gene ontology enrichment using DAVID Functional Annotation Bioinformatics Microarray Analysis v6.8 (Dennis et al., 2003).

### 2.9. Drug-target interaction validation and visualisation

Energy minimization, molecular docking and binding energy calculations were performed to validate the potential drug-target interactions. For this, the crystal structures of the protein targets were obtained from RCSB Protein Data Bank (PDB) (www.rcsb.org) (Berman et al., 2000). The proteins for which no crystal structure is available in PDB was modeled using Phyre2 (Kelley et al., 2015). The refinement of the modeled structures was performed using MOdrefiner (Xu and Zhang, 2011) and GROMACS (GROningen Machine for chemical Simulation) 5.0 package (http://www.gromacs.org/) was used for the energy minimisation using Gromos96 53a6 force field using Steepest descent method.

Non-covalent interactions between the proteins and ligands were assessed using molecular docking studies, for that AutoDock 4.2 (Morris and Huey, 2009) and AutoDock Vina packages (Trott and Olson, 2010) were used. Ligplot+, a graphical system that utilises 3D coordinates to generate the interaction maps was used to visualize the interaction present in protein-ligand complex (Laskowski and Swindells, 2011).

## 3. Results and Discussion

### 3.1. Identification of AEHs and AEDs

We could identify 63 anti-epileptic herbs (AEHs) *via* an extensive review of the PubMed research articles and assigned a unique identifier to each herb. The detailed information of the identified herbs including their scientific name, the unique identifier and the reference source is presented in Table1. A list of 40 anti-epileptic drugs (AEDs) currently used for the management of epilepsy or under clinical trials was also enlisted. The chemical information of these drugs including their DrugBank Id, PubChem Id and SMILES notation is given in supplementary file4, Table S4

### 3.2. Herb-Phytochemical (H-PC) network

The phytochemical (PC) dataset of 63 anti-epileptic herbs consists of unique 867 entries out of total 1993 PCs collected. For the study, each phytochemical of the PC dataset is assigned a Phytochemical-ID (supplementary file5, Table S5). Detailed mapping of these 867 PCs to their corresponding herb is presented in supplementary file6, Table S6. This information was used as an input to construct the H-PC interaction network (Fig. 2). The network analysis shows that several PCs are shared by many of the anti-EP herbs while others are specific in nature. PC157 (palmitic acid) is the most common PC as it is found in 29 out of 63 anti-EP herbs. Other PCs enriched in the AEHs are PC350 (ascorbic acid), PC170 (oleic-acid), PC044 (linoleic-acid) with the degree centrality (C_d_) value of 28, 27 and 25 respectively. PC350 (*i.e*. ascorbic acid) is one of the most widely studied PC having implications in EP (Sawicka-Glazer and Czuczwar, 2014). The deficiency of PC350 is known to contribute in increasing the severity of the PTZ induced seizures (Warner et al., 2015). Out of 867 phytochemicals, 349 were found to have drug-likeliness properties and among these druggable PCs (DPCs); PC703(phosphorus), PC158 (pyridine-3-carboxylic acid) are shared by 24 and 18 anti-EP herbs respectively. Herb “*Zingiber officinale*” (EP57) is found to have a maximum number of DPCs i.e. 104.

**Figure 2.**
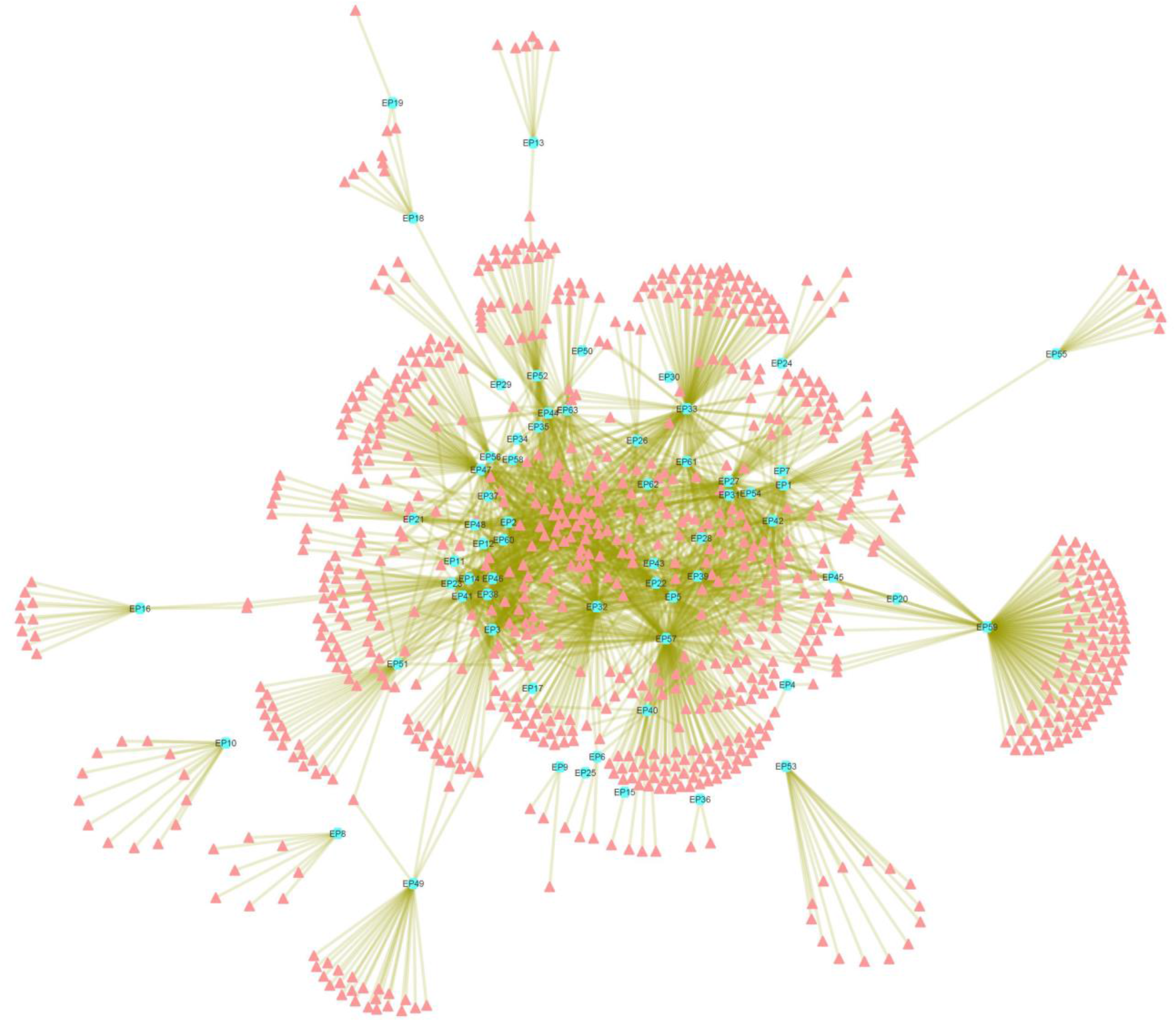
Herb-Phytochemical (H-PC) Network: The network represents associations of 867 unique phytochemicals (peach coloured nodes) with the 63 anti-EP herbs (Cyan nodes). Herb “*Zingiber officinale*” (EP57) is found to have maximum number of phytochemicals among all the anti-EP herbs, followed by “*Piper longum*” (EP50), “*Anethum graveolens*” (EP5) and “*Glycyrrhiza glabra*” (EP33).

### 3.3. Compound classification and clustering

The comprehensive organization of the 349 druggable phytochemicals, obtained from drugability analysis is found to be distributed in the 16 broad chemical classes (Fig. 3A). Detailed classification of the DPCs reveals that the class corresponding to terpenoids especially “sesquiterpenoids” is highly prevalent in this dataset. This observation may be due to the fact that terpenoids constitute the largest and most diverse class of natural compounds (Aharoni et al., 2005; Tholl, 2015). In addition to their role in the plant growth and survival, both primary and secondary metabolites of the terpenoid biosynthetic pathway possess commercial, ecological as well as medicinal significance (Aharoni et al., 2005). Moreover, the dataset is highly enriched for the compounds capable of crossing the blood-brain barrier viz. hydrocarbons, terpenes and carbonyl compounds (Maa and Figi, 2014). Various studies have correlated the anticonvulsant activity of plant-based products with their terpenoids composition (Sayyah et al., 2005; De Sousa et al., 2007). The anticonvulsant properties of sesquiterpenoid metabolites have also been previously identified in zebrafish and mouse seizure models (Orellana-Paucar et al., 2012; Orellana-Paucar et al., 2013). This shows that these compounds may be given further attention for detailed examination as they provide reliable data for the future anti-epileptic drug discovery approaches.

**Figure 3.**
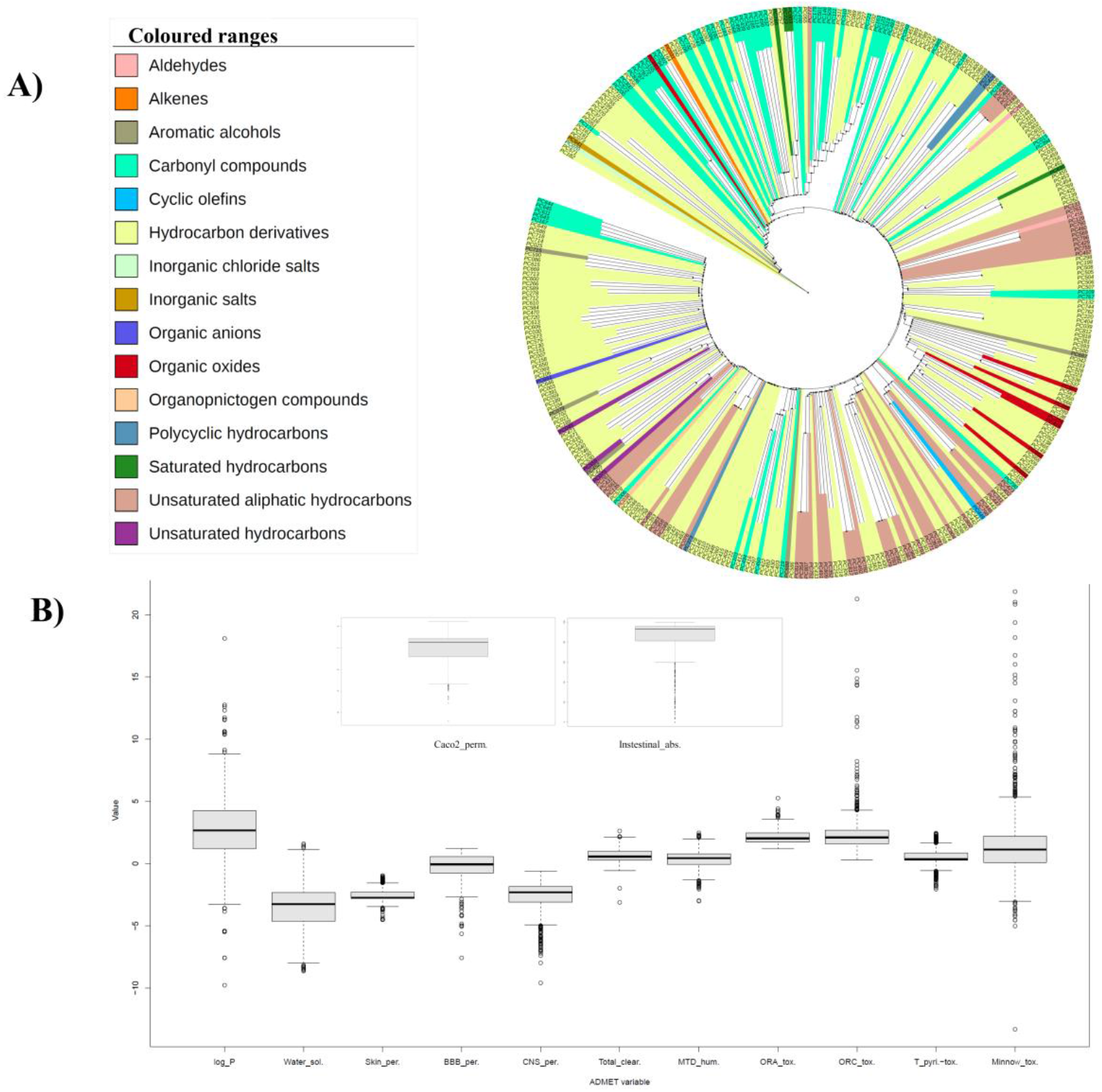
**(A). Clustering and chemical classification of DPCs:** The hierarchical clustering of 348 DPCs based on their atom-pair descriptors and tanimoto coefficient, obtained using Chemmine tool. One DPC corresponding to the PC703 with the chemical class of “Non-metal compounds” could not be clustered and therefore not represented in this tree layout. **(B). ADMET properties of DPCs:** Box and whisker plot represents the 13 ADMET properties of the 349 DPCs. The properties were evaluated using the pkCSM server.

Other than the class of terpenoids, compounds belonging to “Methoxyphenols” are significant in this dataset. Although the use of phenolic compounds in reference to epilepsy is less explored, our findings suggest that the detailed investigation of such compounds could be of considerable importance. The chemical class of each DPC can be checked in supplementary file7, Table S7.

### 3.4. Druggable phytochemical - Protein target (DPC-PT) network

To understand the molecular interactions of DPCs with human proteins, a bipartite DPC-PT network was constructed (Supplementary Fig. 1). This network consisted of the mapping of DPCs (349) with their protein targets (4982) that are obtained using three algorithms of target prediction (described in Materials and Methods). Prioritization of each 16329 phytochemical-protein pairs was performed on the basis of their prediction from each of the three target prediction algorithms. Sixty-six (66) high confidence DPC-PT pairs were identified as these were predicted from all the three target prediction algorithms used (supplementary file8, Table S8).

A sub-network of DPC-PT network [referred as DPC-PTE] (Fig. 4A) consisting of 838 nodes (336-phytochemicals; 502 proteins) and 3002 DPC-PT pairs, specific to the EP-gene pool (mentioned in material and methods section) was derived to highlight the interactions among epilepsy proteins and DPCs that may target them. In this network, three DPCs corresponding to PC116 (glycerol), PC161 (ethanol) and PC043 (17-beta estradiol) possess the target degree centrality value (C_d_) more than 100 *viz*. 259, 140 and 102 respectively. This indicates that these phytochemicals may play a key role in regulating the EP targets *via* regulating multiple proteins simultaneously. From the protein target list, P10636 (C_d_ =95), P10275 (C_d_ =95), P35354 (C_d_ =91) retain the highest number of contacts with the DPCs in DPC-PT network. P10636 is a microtubule-associated protein tau and its involvement in various neurological diseases is well known (Kosik et al., 1986). The microtubule overexpression has been linked to the temporal lobe epilepsy *via* collateral sprouting of hippocampal mossy fibers (Pollard et al., 1994). Also, various AEDs (Anti-epileptic drugs) are shown to induce changes in the expression of androgen receptors P10275, when taken by patients with temporal lobe epilepsy (Killer et al., 2009). Expression of protein P35354 (prostaglandin G/H synthase-2) has been shown to have an elevated induction upon seizures (Marcheselli and Bazan, 1996). To access high confidence interactions of DPC-PTE network, a sub-network corresponding to compound-protein interactions predicted by at least two of the three target prediction algorithms were derived (Fig. 4B). Detailed inquiry of this sub-network helped us to identify the prospective role of apigenin (PC668) in the regulation, based on its protein-targeting capability (C_d_ =9), compared to other DPCs. Among the total 29 DPCs of 66 high confidence DPC-PTE pairs, maximum contribution of 10 from EP33 (*Glycyrrhiza glabra*) and 9 from EP47 (*Punica granatum*) shows that anti-epileptic action of these herbs is worthy of attention.

**Figure 4.**
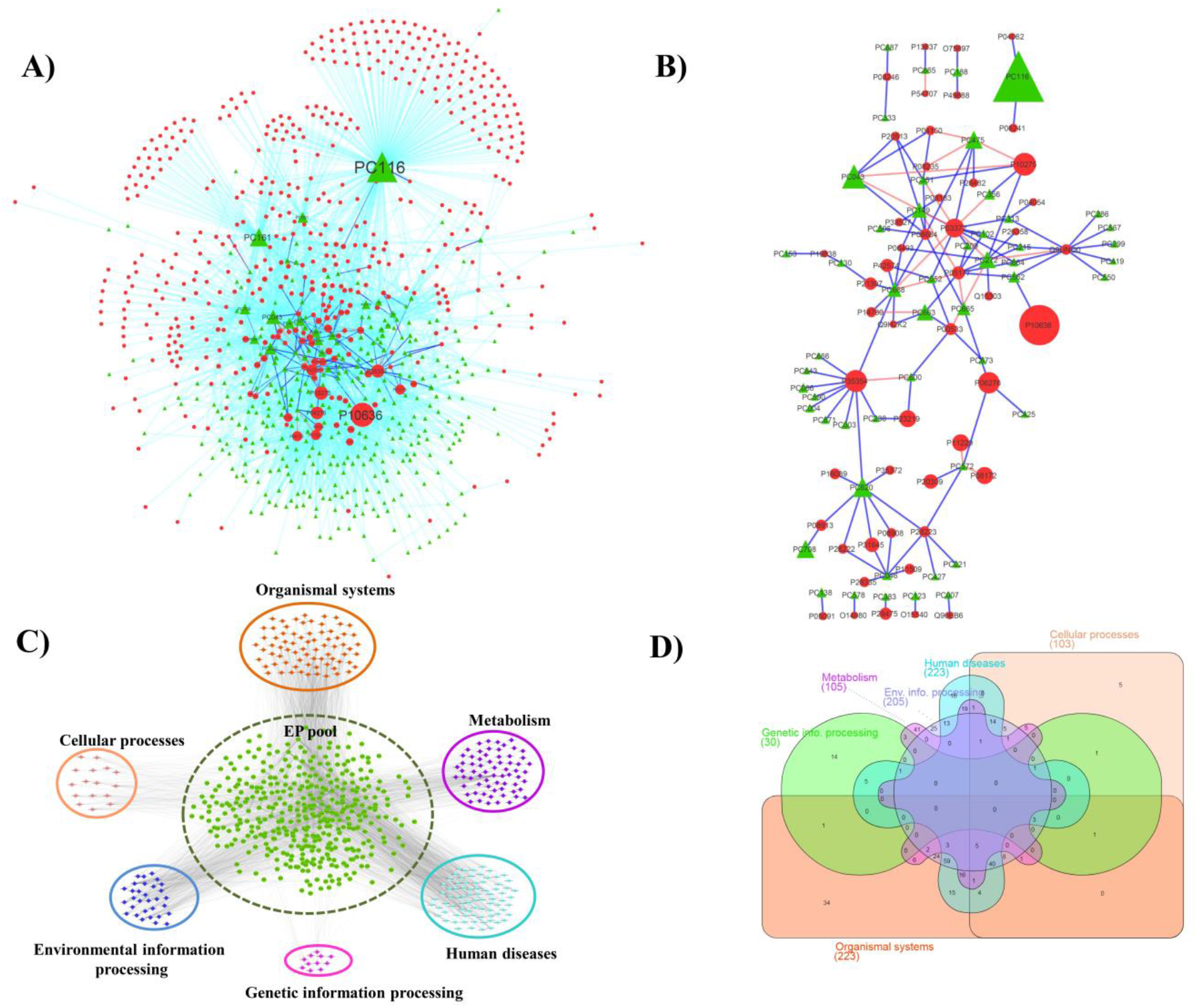
**(A). A sub-network of DPC-PT network specific to Epilepsy (DPC-PTE):** This network consists of 838 nodes (336-druggable phytochemicals; 502 proteins) and 3002 DPC-PT pairs, specific to the EP-pool proteins. Phytochemicals are represented by green coloured nodes and EP-pool proteins with red coloured nodes. Interactions among the phytochemical-protein pairs are either represented as cyan (predicted by any 1 protein target algorithms), blue (predicted by any 2 protein target algorithms) or orange coloured edges (predicted by all 3 protein target algorithms). Size of the nodes is based on their corresponding degree value in the network. Among phytochemicals, PC116 holds the maximum degree value. **(B). A high confidence sub-network of DPC-PTE network:** For specifically examining the high-confidence interactions among the DPCs and protein targets associated with EP, a sub-network of DPC-PTE network is constructed by considering the interactions predicted by either 2 or 3 target prediction algorithms. **(C). PTE-HP (Protein targets associated with Epilepsy - Human pathway) Network:** The pathway enrichment network consists of 677 nodes and 2862 edges, specific to the EP-pool proteins. 400 proteins of EP-pool targeted by any of the DPCs are found to be involved in the 6 broad KEGG pathway classes *i.e*. Organismal systems, Cellular processes, Environmental information processing, Genetic information processing, Human diseases and Metabolism. The 6 main pathway classes are arranged around the proteins of EP-pool, represented in green coloured nodes in the center of the network. **(D). Distribution of EP-pool proteins among KEGG pathway classes:** Venn-diagram representing the distribution of 400 proteins of the PTE-HP network among the 6 broad KEGG pathway classes. The class corresponding to the “Organismal systems” and “Human diseases” are found to be highly enriched with EP-proteins while the class of “Genetic information processing” includes the least number of proteins *i.e*. 30. Many proteins are shared among these classes, the feature attributed towards association of a single protein in multiple pathways.

### 3.5. Protein Target-Human pathway (PT-HP) network

PT-HP network illustrates the association of protein targets (protein targets of DPCs) with human pathways based on KEGG analysis. Detailed mapping of all the 4982 proteins targeted by the DPCs into 314 pathways associated with *Homo sapiens* is presented in supplementary file9, Table S9. Only 3030 proteins out of 4982 protein targets show their involvement in the human pathways, according to KEGG analysis. The data is used as an input to construct a bipartite PT-HP network, consisting of 3344 nodes (3030 protein targets; 314 human pathways) and 15534 edges. Proteins involved in epilepsy were selected from PT-HP network, and a sub-network specific to epileptic proteins and their regulatory pathways was constructed (PTE-HP, Protein Targets of Epilepsy-Human pathway) (Fig. 4C). PTE-HP consisting of 677 nodes and 2862 edges was analyzed for the human pathway enrichment of the EP proteins. This is essential to detect the disease-associated pathways and to understand the underlying mechanism of “protein-disease associations” of epilepsy in detail. Also, the analysis could help in the identification of novel pathways and their regulatory proteins in the context of epilepsy. As shown in Fig. 4C, it is observed that proteins involved in pathways associated with “organismal systems” and “human diseases” are enriched in this dataset. Next in order were the categories belonging to the class of “environment processing” and “metabolism”. Many proteins were shared among these classes, a feature attributed towards association of a single protein in multiple pathways. A set of 9 proteins corresponding to P29475 (C_d_ =11), O60503 (C_d_ =43), P31749 (C_d_ =76), Q9NQ66 (C_d_ =46), P19174 (C_d_ =33), P16885 (C_d_ =31), Q13393 (C_d_ =14), O14939 (C_d_ =12) and P35354 (C_d_ =16) were highlighted in this step, as they are shown to possess multi-level regulatory behaviour *via* its involvement in each of these four major pathway classes. Such proteins may be considered as important targets for the EP because modulation at these levels will give a direct effect on the pathways of their involvement. Distribution of EP proteins among the six major pathway classes is shown in the form of the Venn-diagram constructed using InteractiVenn (http://www.interactivenn.net/)(Heberle et al., 2015) Fig. 4D.

Among the class of “organismal systems”, the pathway corresponding to “path: hsa04724; glutamatergic synapse” share the maximum number of proteins. Regulation of glutamate homeostasis plays a key role in the pathophysiology of epilepsy (Coulter and Eid, 2012) and altered behavior of this pathway upon the episode of neonatal seizures has also been reported (Cornejo et al., 2007). Also, the “path: hsa01100” (metabolic pathways) and “path: hsa04080” (Neuroactive-ligand-receptor interaction pathway) have a maximum number of EP proteins involved in them. These findings support previous studies according to which there exists a therapeutic relationship between brain activity and metabolism to treat neurological disorders (Masino et al., 2009)(Ruskin and Masino, 2012). Although direct correlation of the “path: hsa04080” and epilepsy could not be established, but this pathway has remained as an important source for the development of various therapeutic strategies against various neurological disorders (Kong et al., 2015). Thus, it may be correlated that regulation of genes corresponding to this neurological pathway may be responsible for the wide spectrum alteration in various neurological functions in epilepsy. Detailed information of human pathways with the number of EP pool proteins associated with them, is presented in supplementary file10, Table S10.

In the following, we detail our network analysis for some specific case studies.

#### Case study-I: Investigation of the neuromodulatory effects of AEHs

To explore the neuromodulatory potential of AEHs, we attempted to uncover the underlying mechanism of neuromodulation by highlighting the potential DPC-PT interactions potentially responsible for their regulation. Towards this end, pathways specific to nervous system and protein targets involved in these pathways were selected for the further analysis. A sub-network specific to all the 10 KEGG pathways associated with nervous system namely, glutamatergic synapse (path:hsa04724), GABAergic synapse (path:hsa04727), cholinergic synapse (path:hsa4725), dopaminergic synapse (path:hsa4728), serotonergic synapse (path:hsa4726), long-term potentiation (path:hsa04720), long-term depression (path:hsa4730), retrograde endocannabinoid signaling (path:hsa04723), synaptic vesicle cycle (path:hsa04721) and neurotrophin signaling pathway (path:hsa4722) and their associated proteins was focused in detail. This sub-network consisting of 110 nodes (100 proteins; 10 pathways) and 200 edges was analysed in detail for the following sub-cases (Supplementary Fig. 2):

#### Ia: Identification of regulatory DPCs for metabotropic glutamate receptors (mGluRs)

The pathway relationship of above sub-network shows that the glutamatergic synapse (path:hsa04724) is associated with 32% of the protein targets and retrograde endocannabinoid signaling (path:hsa04723) is associated with 27% of the protein targets. Mapping of the protein targets in the glutamatergic synapse pathway (path:hsa04724) is shown in Fig. 6A. As already discussed about the role of Glutamatergic mechanism in the condition of seizures and epilepsy, detailed literature data could link up the role of metabotropic glutamate receptors (mGluRs) for inducing epileptic symptoms in the patients (Barker-Haliski and Steve White, 2015; Huberfeld et al., 2011). Also, many of the current antiseizure drug development strategies are focused on the use of compounds that can potentially modulate the glutamatergic signaling *via* mGluRs (Barker-Haliski and Steve White, 2015). For the identification of such potential drug candidates from our dataset of AEHs, proteins corresponding to mGluRs were backmapped in the DPC-PT network. This led to the selection of 11 DPCs corresponding to (PC146; caproic acid), (PC161; ethanol), (PC163; lauric acid), (PC179; butyric acid), (PC461; camphor), (PC496; propionic acid), (PC501; valeric acid), (PC620; melatonin), (PC650; benzoic acid), (PC708; lactic acid) and (PC743; 2-nonynoic acid). A sub-network corresponding to the interactions between 11 DPCs and 8 proteins (corresponding to mGluRs), presented in Fig. 6B was evaluated for examining the multi-targeting nature of these DPCs. We found that PC461, PC708 and PC620 target 6 out of 8 mGluRs. Docking study was carried out for each of these interaction pairs and binding free energy values were also calculated that are mentioned as the edge weights in the network shown in Fig. 6b. The best docked conformation was selected based on the binding energy value, *i.e*. one with lowest binding energy and was used to represent the molecular interactions via Ligplots. The DPCs are shown to possess a very good binding affinity to the mGLuRs in the range of −3.1 to −7.3 kcal/mol. The best binding energy was observed for PC620 with mGluR3, mGluR6 and mGluR8. The 2D representation of the interactions shows that the targets of PC620 are involved in multiple hydrogen bonds and hydrophobic contacts (Fig. 6C).

Further mGluR5 (P41594) protein was checked for its regulatory phytochemicals and 8 DPCs (PC146, PC161, PC163, PC179, PC496, PC501, PC650 and PC743) could be selected in this regard (see Fig. 6B). We selected this protein for the detailed analysis because mGluR5 antagonists were earlier reported to be effective in the management of seizures (Hagerman et al., 2009; Youssef et al., 2018). Such antagonists may be looked upon to control the seizures in case of epilepsy also. The above mentioned 8 DPCs were mapped in H-PC network to identify the AEHs containing them. Mapping of 5 of these 8 DPCs in EP-42 (*Ocimum gratissimum*) and 3 in both EP-33 (*Glycyrrhiza glabra*) and EP-26 (*Croton tiglium*) reflect their mGluR5 regulatory behavior. It was interesting to note that the selected 8 regulators of mGluR5 also show their affinity towards mGluR1. The binding affinity of each of the 8 DPCs with mGluR5 (P41594) and mGluR1 (Q12355) are represented along the edges of the network shown in Fig. 6B. Detailed examination of the interaction pairs and their associated binding affinity draws attention towards the multi-targeting and synergistic behavior of DPCs (see Fig. 6b). While PC620 is shown to hold a multi-targeting property for mGluR3 and mGluR6 at a good binding affinity score, synergistic action of PC146 and PC501 to regulate mGluR5 is also significant. The interaction analysis also supports the multi-targeting and synergistic action of DPCs (Fig. 6C). The data can be investigated in detail, for the screening of lead molecules that may be found effective for their anti-epileptic actions.

**Figure 6.**
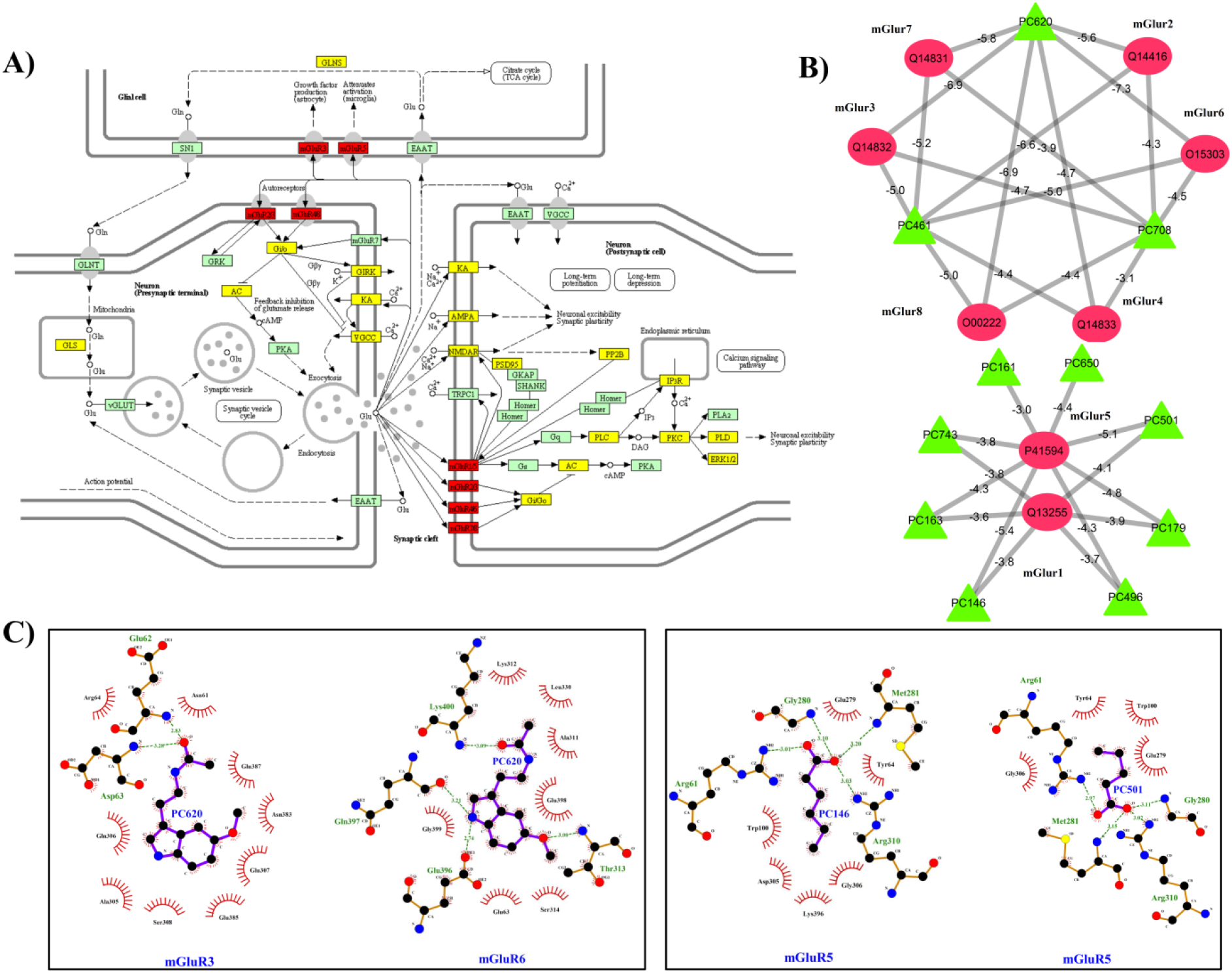
**(A). Glutamatergic synapse (path:hsa04724) pathway mapping of the protein targets of DPCs:** The location of protein targets of DPCs in the KEGG pathway hsa04724 is represented in yellow and red coloured boxes, red ones are specific to mGluRs. **(B). Sub-network of mGluRs and their regulatory DPCs:** Green coloured nodes of the network correspond to 11 DPCs (PC146, PC161, PC163, PC179, PC461, PC496, PC501, PC620, PC650, PC708 and PC743) that possess the tendency to regulate 8 classes of mGluRs (Red coloured circular nodes). The binding energy values (kcal/mol) of the mGluRs and their regulatory DPCs are represented along with their corresponding edges in the network. For docking studies, the structure of mGluR6, mGluR4 and mGluR8 were modelled using PHYRE2 while for others they were obtained from RCSB-PDB with following PDB-IDs: 3KS9 (mGluR1), 4XAS (mGluR2), 6B7H (mGluR3), 3LMK (mGluR5) and 3MQ4 (mGluR7).**(C). The interaction analysis of the mGluRs and their regulatory DPCs:** Docked complexes with the best binding energy values are analysed for the hydrophobic interactions as well as hydrogen bonded residues using Ligplot+. The ligplots enclosed in boxes are the representative cases of multi-targeting and synergistic actions of selected phytochemicals respectively. Protein residues represented along the arcs are involved in the hydrophobic interactions whereas the residues involved in the hydrogen bonding are represented using dashed lines. The PC620 is shown to possess much negative binding energy compared to the other 10 DPCs and the interaction analysis highlights the importance of the acetamide group (-NHCOCH_3_) in the interaction.

#### Ib: Screening of novel neuromodulatory DPCs

Towards prioritization of DPCs obtained from AEHs against the above mentioned 100 proteins involved in neuromodulatory pathways, we mapped these proteins to the ‘approved protein target list’ provided by DrugBank. 81 proteins could be successfully mapped in this approved list and the further study was restricted to the set of these 81 neuromodulatory proteins only (NM-proteins). For exploring the potential of DPCs in regulating the NM-proteins, these proteins were backmapped to the DPC-PT network and DrugBank database. From the DPC-PT network, 241 DPCs were found to have a role in the regulation of NM-proteins. From the DrugBank database, 1684 drugs (of total 2337 approved drugs listed in DrugBank, **D dataset**) were found to target 1311 proteins (of 4982 PTs of all DPCs). Out of these, a total of 467 drugs were found to be associated with the selected 81 NM-proteins. In this manner the regulatory molecules of the selected 81 NM-proteins were obtained from two facets, one which includes the DPCs of anti-EP herbs (241 DPCs, **D1 dataset**) and the second includes drugs given in DrugBank database (467 drugs, **D2 dataset**). The drugs and DPCs of D, D1 and D2 datasets with their associated proteins are given in supplementary file11, Table S11.

“Similar compounds tend to generate similar biological effect” is the key concept that forms the basis of medicinal chemistry (Martin et al., 2002). Applying this idea in our study, we attempted to screen potential phytochemicals that may produce similar effects to the already existing drugs. We first calculated the pairwise Tanimoto coefficient (TC) values of 349 DPCs against all the 2159 drugs related to “drugbank_approved_structure” dataset of DrugBank. TC similarity values are presented in form of heatmap (Fig. 5). A subset of TC values of 241 DPCs (D1 dataset) with the approved drugs present in DrugBank (1684 **drugs, D dataset**), is selected for further analysis. To identify novel molecules from the DPCs that may have the potential to exhibit similar effects as of the existing drugs, selection criteria comprising two similarity conditions was applied to all the pairs. A DPC was considered for the further analysis, only if (i) it has TC value >0.85 with any of the drug of D dataset (Gamo et al., 2010), (ii) it should not have the TC value =1 and also should not have SMILES identical with any of the drug in the D dataset.

**Figure 5.**
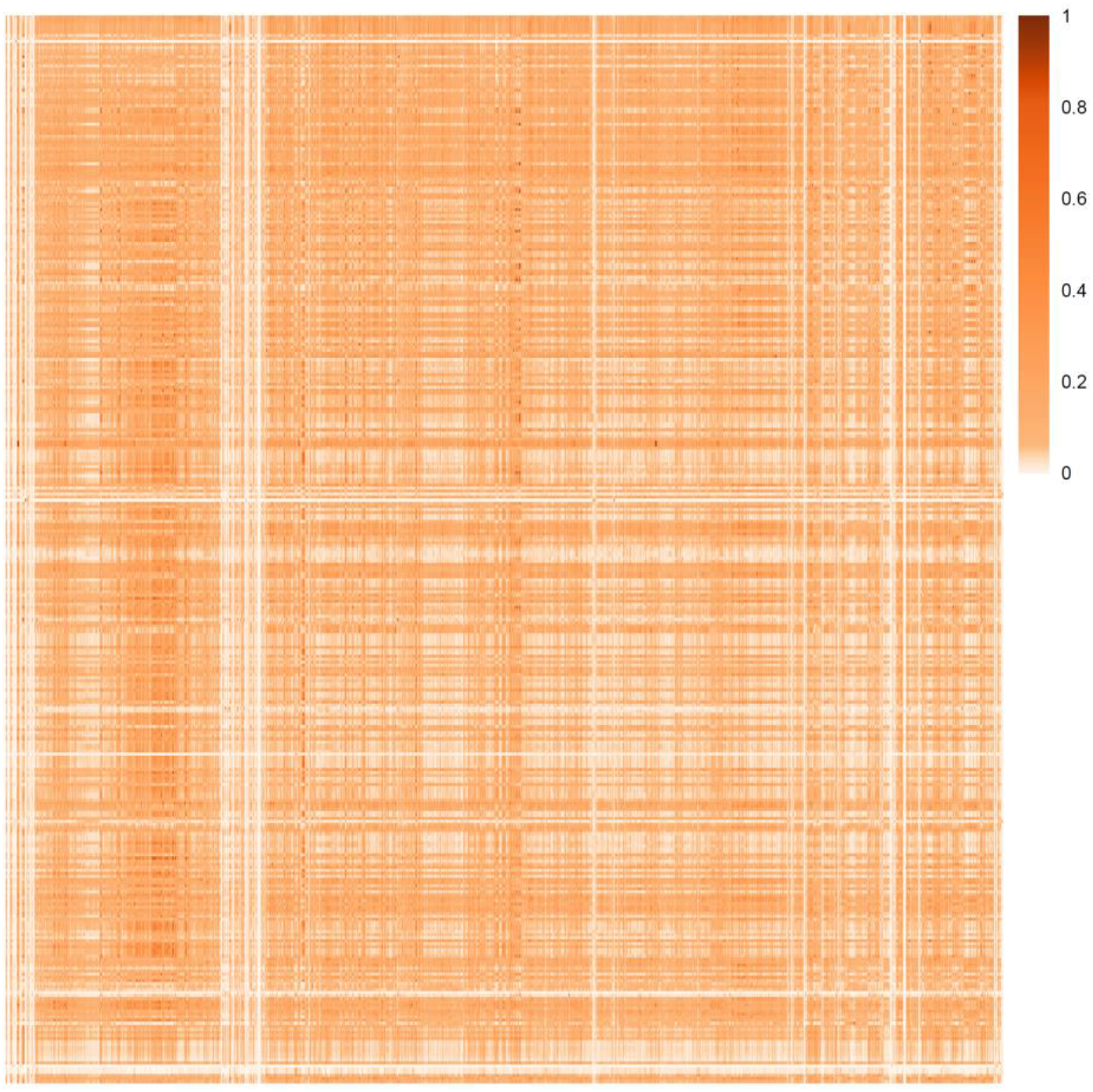
Heat-map corresponding to the Tanimoto coefficient based similarity of the DPCs with the drugs of the DrugBank: The heat-map corresponds to the molecular similarity of 349 DPCs and 2159 drugs related to “drugbank_approved_structure” dataset, based on the Tanimoto score. The columns of the matrix represent DPCs and rows represent drugs.

Satisfying this selection criterion, 11 DPCs (PC179, PC215, PC427, PC501, PC590, PC613, PC668, PC671 and PC663) were screened-in against 188 DrugBank compounds and these two sets of compounds are found to be connected via 13 NM proteins. A tripartite network highlighting the interactions of 13 NM proteins with their corresponding 11 DPCs and 188 drugs shows that the multi-targeting effect of PC668 is noteworthy. As seen in Fig. 7, PC668 holds the potential to regulate 11NM proteins of the network, which include three high confidence interactions with the P35354, P21397 and P42574. In addition to this, PC427, PC590 and PC67 also constitute the interaction partner of high-confidence pairs. Since these 11 DPCs maintain the drugability and similarity criteria used in this study, we propose these 11 DPCs for the detailed investigations by exploring their efficacies towards regulation of target protein activities *via in-vitro* and *in-vivo* experiments with a special focus on PC668, PC427, PC590 and PC67. The concept of designing synthetic analogues could also be explored to obtain satisfactory therapeutic effects from these DPCs.

**Figure 7.**
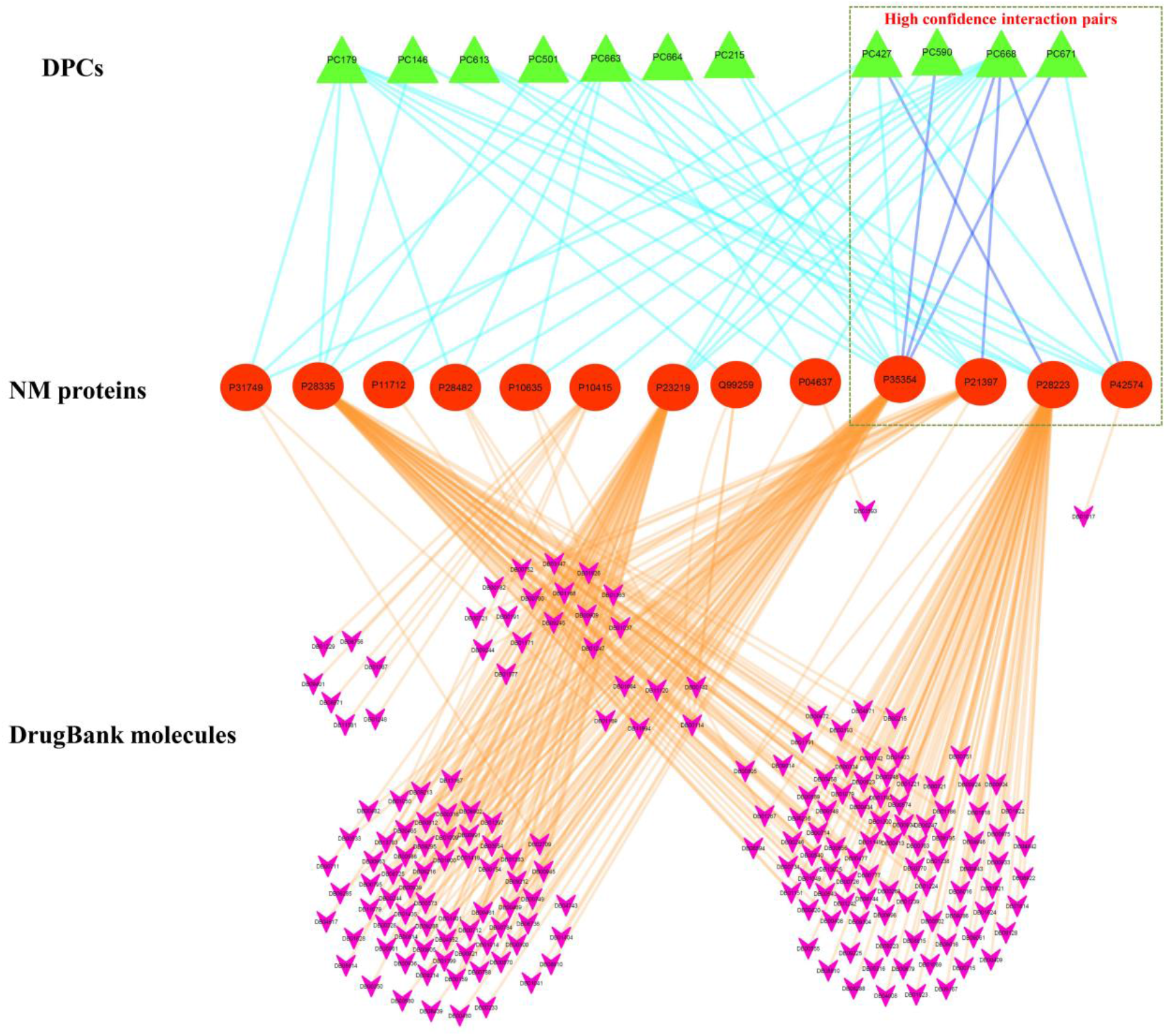
Tripartite network of NM proteins with their interacting DPCs and DrugBank molecules: The network consists of 212 nodes and 347 edges, consisting of interactions of 13 NM proteins (represented in the middle layer; red coloured nodes) with the 11DPCs (represented in the top layer; green coloured nodes) and 188 drugs of the DrugBank (represented in the bottom layer; pink coloured nodes). Edges of the network are coloured differentially just for making distinctions among the interactions of DPCs and drugs with NM proteins. The edges representing interactions among DPCs and NM proteins are either represented in cyan (predicted by any 1 protein target algorithms) or blue (predicted by any 2 protein target algorithms). Six high confidence interaction pairs were identified in the network, consisting of the interaction of 4 DPCs with 4 NM proteins are represented in a triangular box. No interaction pair predicted by all the methods appeared in the network. Interactions of the NM proteins with their associated drugs are represented with orange coloured edges.

#### Case study-II: Searching DPCs having poly-pharmacological similarity with AEDs

For the screening of DPCs that may have a tendency to elicit similar effects compared to the AEDs, a comparative analysis of the protein targets of both DPCs and AEDs was carried out. 40 AEDs were found to interact with 1045 human proteins, forming 1747 drug-protein pairs (listed in supplementary file12, Table S12). To recall, 349 DPCs were found to interact with 4982 proteins. In the further analysis, we considered compound-protein interactions predicted by at least two of the three target prediction algorithms only. 607 interactions from DPCs and 161 from AEDs qualify this criterion.

35 proteins were common in both the datasets that were analysed further by constructing a tripartite network, to highlight their regulators from both DPCs and AEDs lists. AED19 and AED04 both are found to have interactions with three protein targets, namely P11511, P10275 and P11413. Association of these proteins is well studied in either direct correlation with epilepsy or with the onset of the seizure during epilepsy. When looked into the network, it is observed that the PC043 is a DPC that can regulate all the three protein targets of AED19 and AED04 (Fig. 8). Also to note, all of these three interactions of PC043 have been reported by all the three target prediction algorithms.

**Figure 8.**
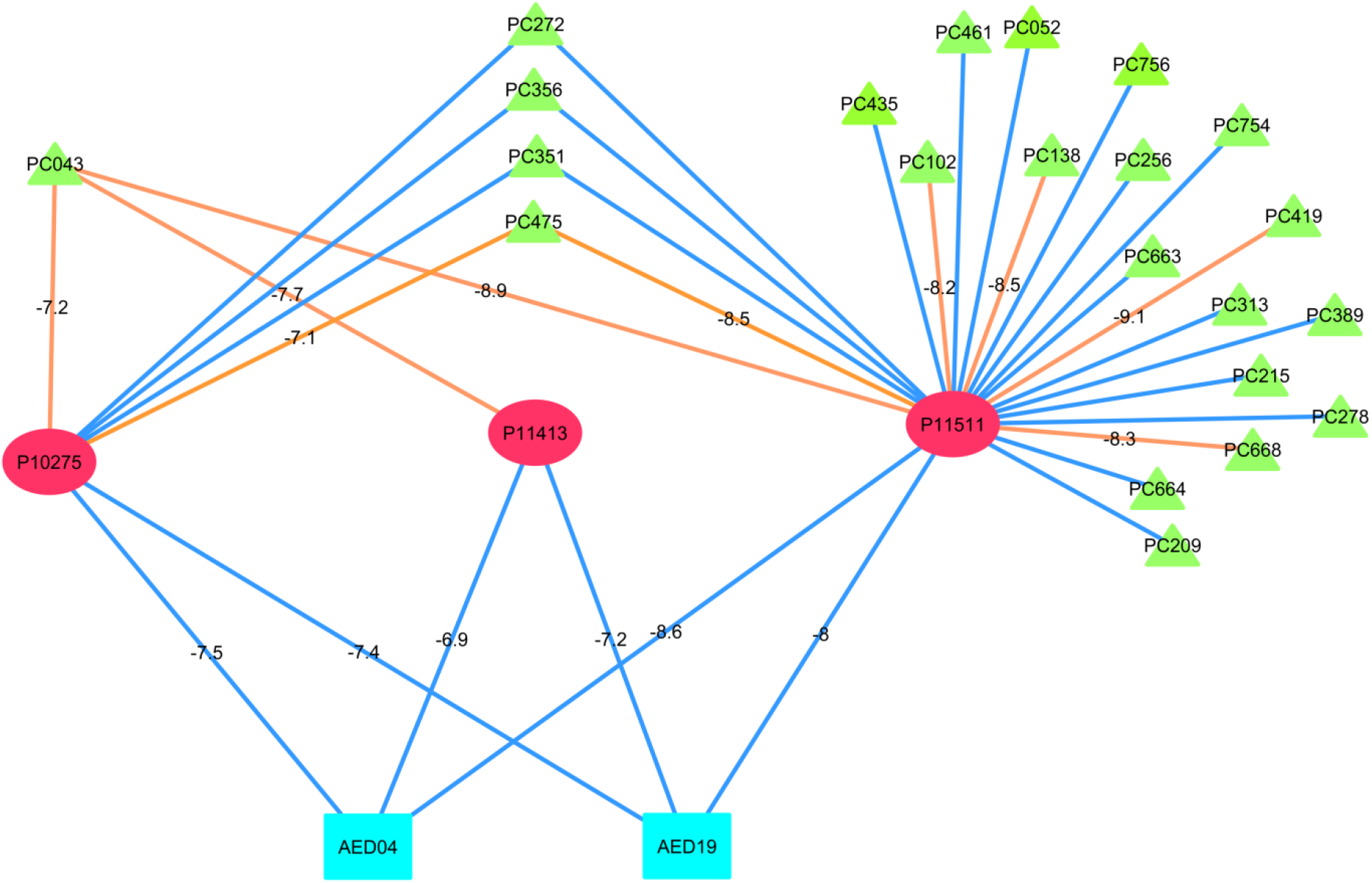
Multi-targeting potential of DPCs to regulate protein targets of AEDs: The tripartite network represents the multi-targeting potential of DPCs to regulate the protein targets of AEDs. Three protein targets (P10275, P11413 and P11511) commonly targeted by 22 DPCs (Green triangular nodes) and 2 AEDs (cyan rectangular nodes) are represented in the middle layer of this tripartite network. The interactions of DPCs and AEDs with their protein targets are either represented as blue (predicted by any 2 protein target algorithms) or orange coloured edge (predicted by all 3 protein target algorithms). The binding energy values (kcal/mol) of the protein targets and their regulatory DPCs and AEDS are represented along with their corresponding edge in the network. Binding energy values show that these DPCs possess a good binding affinity for the protein targets of AEDs, in the range of −7.1 to −9.1kcal/mol. PC043 is shown to target all the 3 proteins with binding energy values comparable to their corresponding AEDs, even better for few cases like AED19 and PC043 for P11413. For docking studies the following PDB IDs of proteins were used;1XOW (P10275),1QKI (P11413) and 3EQM (P11511). Here, only binding energy calculations corresponding to high confidence DPC-PT interactions (i.e. predicted by 3 protein target algorithms) were considered.

P11511, an aromatase protein is a key enzyme of estrogen biosynthesis pathway. The role of aromatase inhibitors as a useful adjunct in the seizure control shows that the screening of regulatory DPCs for P11531 could be of significant importance in treatment procedures of EP, especially in male candidates (Harden and MacLusky, 2005; Harden and MacLusky, 2004). Androgen receptor (AR), P10275 is highly abundant in several regions of the brain and the consequence of their activation by various androgens, like testosterone and its metabolites are responsible for the regulation of behavior and other neuronal functions. As earlier mentioned, changes in the expression of ARs by AEDs in the case of patients with temporal lobe EP shows that the modulation of ARs holds a great potential in disease management (Killer et al., 2009). The role of P11413 (Glucose-6-phosphate 1-dehydrogenase-an important enzyme responsible for glucose metabolism) in EP is based on the correlation of energy depletion and seizure development. Impairment in the glucose metabolism have been examined in various epileptic patients and other epilepsy-related models (McDonald et al., 2018).

#### Case study III: Exploring the potential of DPCs to regulate a functional module in the EP-PPI network

To investigate the comprehensive effects of EP proteins on the entire human PPI, a sub-PPI regulated by EP proteins was constructed and analysed. For that, a high confidence human PPI network was constructed using a STRING score of ≥ 0.9 in which a total of 794 EP proteins could be mapped. Out of these, 611 EP proteins were showing their interactions with other EP proteins (*i.e*. these 611 own the tendency to regulate each other) while remaining existed as individual nodes. Isolated nodes were removed from the network and the resulting network consisting of 611 nodes and 2,681 edges (referred to as **EP-PPI network**) was considered for the further analysis (Fig. 9A). The network degree distribution shows a power-law with *y* = 2 38.92*x*^−1.31^ (Fig. 9b).

To get an insight of the overall organization and the inter-relationship of EP proteins in the EP-PPI network, this network was subjected for modularity analysis using MCODE algorithm that returned 21 densely connected regions *i.e*. clusters. To assess the biological role of each identified cluster, gene ontology (GO) based enrichment of the genes associated with various clusters were examined using DAVID. The complete description of the identified clusters, proteins associated with them and their GO-based annotation are given in supplementary file13, Table S13.

As shown in Table S9, the clusters are found to be involved in multiple biological processes, majority being associated with metabolism and signaling events. Proteins of modules 3, 6, 7 and 9 were directly or indirectly linked with the metabolic processes. For example, module-6 includes one of the major key players of the epileptogenesis, i.e. MMP9 (P14780) gene that has a special role in temporal lobe epilepsy (Wilczynski et al., 2008) and also in the cases of intractable epilepsy (Konopka et al., 2013). In order to gain the information of regulatory potential of MMP9 with respect to other EP proteins, position of this node was checked in both EP-PPI network and its corresponding module. In EP-PPI, its C_d_ value is 20 implying that MMP9 can regulate 19 other EP proteins of this network, while in its respective module (module-6) it shows direct links with 3 of these 19 proteins. This protein was also assessed for an important network measure known as betweenness centrality (C_b_), which reflects the frequency of a node to lie on the shortest path with the other nodes of the network (Hahn and Kern, 2005). Network analysis shows that P14780 holds the maximum C_b_ value of 0.53 among all the other proteins of module-6, thereby reflecting the key relevance of this protein in terms of its information spreading capacity. Discerning the regulatory importance of MMP9 in this module, we found it essential to identify the DPCs that have potential to regulate the activities of MMP9 because the protein lies in the center of this module and regulation at this level will tend to generate global effects on the entire module. For detailed investigation, a sub-network corresponding to module-6 and its regulatory DPCs was constructed (Fig. 9C). As shown, 22 DPCs are having interactions with the proteins of this module, amongst those 13 DPCs are multi-targeting that interact with MMP9 and other proteins also. DPCs corresponding to PC663 (luteolin) and PC668 (apigenin) are found to possess multi-targeting capacity at a high confidence score, thereby indicating a scope for their future investigation. In addition, roles of PC272 and PC043 may also be explored in detail as both of them are found to directly regulate 4 proteins of the module. Using the similar approach other key proteins and their regulators at the module level could be identified and studied in detail.

In the following, we briefly discuss the proteins involved in other modules that are enriched in metabolic processes (that are 3, 7 and 9). Proteins of module-7 are generally associated with folic acid metabolism. The significance of the folic acid in the health of the epileptic patients is basically due to the various vascular and neurological conditions that appear in its deficient state (Moore, 2005). Module-3 is rich in genes associated with the cytochrome P450 family like CYP2C9, CYP3A4, CYP2E1 etc. A significant increase in the expression of these genes like CYP2E1 has been observed in the conditions of spontaneous recurrent seizures (Runtz et al., 2018). Also, pharmacogenetic effects of cytochrome P450 on various disease states, especially CYP2C9 in epilepsy have been established in previous studies (Ingelman-Sundberg, 2004). The role of proteins of module 9 in epilepsy is established based on various studies according to which regulation of glutamate homeostasis is of key importance in the disease process. Glutamate is a prominent excitatory amino acid and impaired transport system of the same has been observed in the condition of epilepsy (Molinari et al., 2005). Further glutamate-astrocyte system, in maintaining glutamate homeostasis is known to play an important role in disease pathogenesis (Coulter and Eid, 2012).

Proteins of module-1 and module-8 are majorly involved in G-protein coupled receptor signaling pathway. These clusters were highly enriched in genes like GRM (glutamate metabotropic receptors), NPY (neuropeptide Y receptors) and HTR (hydroxytryptamine receptors). NPY receptors are densely located in the region of strata radiatum of brain and their altered expression in epileptic brain tissues have been noted earlier (Vezzani and Sperk, 2004). Role of HTR receptors in epilepsy is mainly through their participation in the serotonergic neurotransmission process. Other signaling events associated proteins are from module 4, 5 and 11 corresponding to GO related to gamma-butyric acid signaling, MAPK cascade and signal transduction respectively. Gamma butyric acid (GABA) is one of the important and most widely studied classes of neurotransmitters in relation to epilepsy. GABA receptors are the well-known markers for the disease and the role of GABAergic transmission in epileptogenesis is well studied (Aroniadou-Anderjaska et al., 2008). Since the underlying mechanisms for the disease are diverse in nature, therefore a single event is not considered responsible for disease development. Recently, the roles of MAPK targets have been successfully proved in regulating the synaptic excitability that appears in epileptic conditions (Pernice et al., 2016).

Since the underlying mechanisms for governing the onset of epilepsy are diverse in nature, a similar trend has been observed at the modular analysis of the EP-PPI. Though the proteins associated with epilepsy are involved in various biological processes based on GO terms, majority of them are focused on the metabolic and signaling events of the cell.

**Figure 9.**
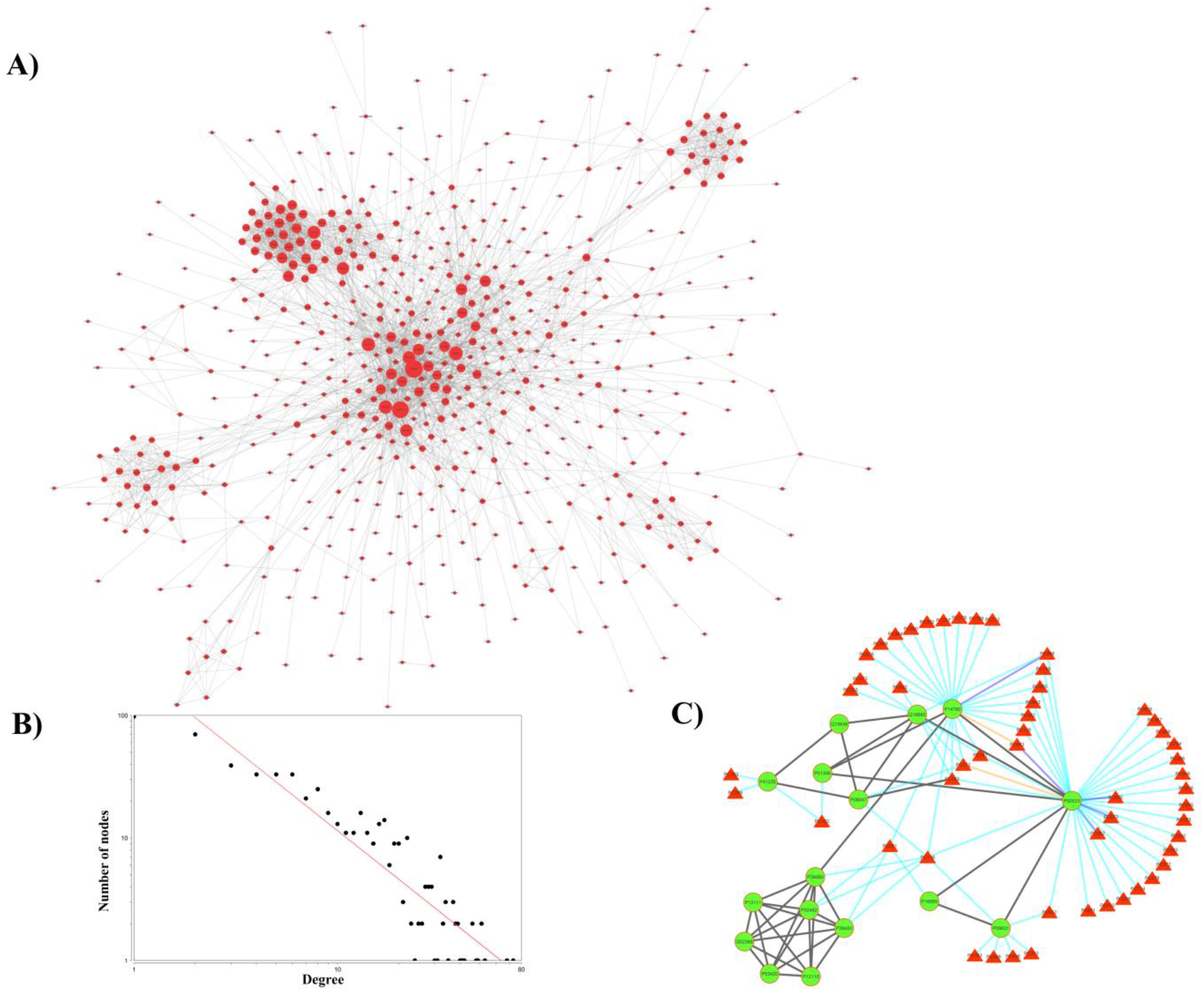
**(A). EP-PPI network:** The network (556 EP-pool proteins with 2639 interactions among themselves) represents the sub-network of human PPI, obtained at the STRING confidence score of > =900. The size of the nodes is based on their corresponding degree value in the network. **(B). Node degree distribution graph of EP-PPI network:** The connectivity of each node of EP-PPI network is represented using node-degree distribution graph. X-axis represents the degree value and Y-axis represents the corresponding number of nodes. Values at both the axes are plotted on a log scale. Network follows a power-law degree distribution with *y* = 238.92*x*^−1.31^. **(C). Interactions among the proteins of module 6 and their regulatory phytochemicals:** The 16 proteins of module 6 are represented as green nodes while the interactions among them are represented by black coloured edges. P14780 corresponding to gene MMP9 is directly known to regulate 3 other proteins of this module. The regulatory phytochemicals of proteins are represented as red coloured triangular nodes and added to the module from PC-PT network. The interactions among phytochemical-protein pairs are either represented as cyan (prediction from any 1 target prediction algorithms), blue (prediction from any 2 target prediction algorithms) or orange coloured edges (prediction from all 3 target prediction algorithms).

## 4. Summary

Despite the recent developments in preclinical research that have facilitated the scientific community to discover valuable and effective drugs against epilepsy, occurrence of the seizures is neither accurately predictable nor under control in more than one-third of the patients affected by epilepsy. Traditional Indian medicinal system has always remained a great source of information about the implications of various herbs and herbal formulae for the management of various diseases. Though the usage of herbal medicines in epileptic treatment is well known for several thousand years, lack of robust confirmation regarding their efficacy has limited their usage in the present scenario. Therefore in this study, we have examined the anti-epileptic potential of various traditional Indian herbs using network pharmacology approach towards discovery of novel drug-like molecules. An extensive search about the anti-epileptic herbs prescribed in Ayurveda resulted in the identification of 63 herbs; detailed information about the phytochemicals content in these herbs forms the basis of present study. To reveal component level anti-epileptic effects of the selected herbs, these were subjected to database screening for identification of their phytochemical composition. Out of total 1993 phytochemicals collected, 867 unique entries were identified based on their chemical information. 349 phytochemicals from this list of 867 are found to have the drug-like properties and are reported as putative drug-like phytochemicals (DPCs). The polypharmacological effects of DPCs are evaluated on the basis of their human-protein targeting capability. To limit the phytochemical-protein interactions specific to epilepsy, an epilepsy gene pool comprising of 1,179 genes (EP-pool) is constructed. Only 336 DPCs show the potential to interact with the proteins of EP-pool. P10636 (i.e. microtubule-associated protein tau) is highlighted as one of the majorly targeted proteins by the DPCs (95 in number) of AEHs. Eleven potential regulators of mGluRs with their supporting docking and interaction analyses are highlighted in this work. Among these, 3 DPCs *i.e*. PC146 (caproic acid), PC501 (valeric acid) and PC179 (butyric acid) also show similarity with DrugBank molecules,thereby indicating their possible usage in mGluR5 and mGluR1 regulators design strategies. Role of EP33 (*Glycyrrhiza glabra*) and EP47 (*Punica granatum*) as the major sources of potential drug-like phytochemicals having mGluRs regulatory potential in epilepsy is explained. Four DPCs namely, PC663 (luteolin), PC664 (pinocembrin), PC251 (pectin) and PC668 (apigenin) that successfully passed through the DrugBank molecules similarity criterion have shown the propensity to interact with all the protein targets of two AEDs. Among the above mentioned 4 DPCs, PC663 and PC668 are found to regulate the module-6 of EP-PPI network either by directly interacting with the protein P14780 (having highest C_b_ value) or its neighbors. The presence of high confidence interactions of PC668 (apigenin) in both the case studies corresponding to DrugBank molecules and AEDs, establishes a strong regulatory potential of this DPC against epilepsy. The presence of this DPC in both EP33 and EP47 further strengthen the anti-epileptic action of these herbs. The multi-therapeutic effects elicited by PC043 (17-beta estradiol) are predicted as it is found to simultaneous interact with all the 3 proteins targets of AEDs. It is interesting to note that some DPCs, when analysed for their binding energy values show much better negative binding energy values for the protein-targets compared to the corresponding AEDs.

Comparison of the screened DPCs having multi-targeting and/or synergistic effects with the existing drugs of DrugBank and currently used anti-epileptic drugs performed in this study will be a substantially important resource for future anti-epileptic drug design strategies. Also, the representation and analysis of complex DPC-PTE interactions and DPC-Pathway associations in the form of network maps will be significantly useful to the researchers working in this area to further explore this deluge of information in a relatively simple mode. We believe that hierarchy of steps used in this computational framework will be immensely helpful to delineate the phytochemical-level anti-epileptic potential of the herbs of not only Indian Ayurvedic system but other traditional medicinal systems of the world also in a much clearer, detailed and meaningful manner against any class of disease or disorder. The study may be considered as a major step towards integrating the wealth of traditional knowledge with scientific outlook for their application in modern day therapeutics to meet the current demand of drug-discovery.

## Acknowledgments

N.C. is grateful to the Indian Council of Medical Research (ICMR) for support provided through ICMR-SRF. N.C. and V.S. thank the support of Central University of Himachal Pradesh for providing computational facilities.

## Abbreviations

ADMET: Absorption, Distribution, Metabolism, Excretion and Toxicity
AEDs: Anti-Epileptic Drugs
AEHs: Anti-Epileptic Herbs
CAMs: Complementary and Alternative Medicines
DALYs: Disability Adjusted Life Years
DPC-PT: Druggable PhytoChemical-Protein Target
DPC-PT-BP: Druggable PhytoChemical-Protein Target-Biological Pathway
DPC-PTE: Druggable Phytochemical-Protein Targets associated with Epilepsy
DPCs: Druggable PhytoChemicals
EP: Epilepsy
EP-PPI: Epilepsy-Protein Protein Interaction
GO: Gene-Ontology
GROMACS: GROningen MAchine for Chemical Simulation
H-PC: Herb-Phytochemical
KEGG: Kyoto Encyclopedia of Genes and Genome
MCODE: Molecular COmplex Detection
mGluRs: metabotrophic Glutamate Receptors
NM-proteins: NeuroModulatory-proteins
PCs: PhytoChemicals
PDB: Protein Data Bank
PT-HP: Protein Target-Human Pathway
TC: Tanimoto Coefficient

